# *De novo* assembly and characterization of transcriptome towards understanding molecular mechanism associated with MYMIV-resistance in *Vigna mungo* - A computational study

**DOI:** 10.1101/844639

**Authors:** Manoj Kumar Gupta, Ravindra Donde, Gayatri Gouda, Ramakrishna Vadde, Lambodar Behera

## Abstract

The fast climate change affects yield in *Vigna mungo* via enhancing both biotic and abiotic stresses. Out of all factors, the yellow mosaic disease has the most damaging effect. However, due to lack of reference genome of *Vigna mungo, the* complete mechanism associated with MYMIV (Mungbean Yellow Mosaic Indian Virus) resistance in *Vigna mungo* remain elusive to date. Considering this, the authors made an attempt to release new transcriptome and its annotation by employing computational approaches. Quality assessment of the generated transcriptomes reveals that it successfully aligned with 99.03% of the raw reads and hence can be employed for future research. Functional annotation of the transcriptome reveals that 31% and ∼14% of the total transcripts encode lncRNAs and protein-coding sequences, respectively. Further, analysis reveals that, out of total transcripts, only 4536 and 78808 are significantly down and up-regulated during MYMIV infection in *Vigna mungo*, respectively. These significant transcripts are mainly associated with ribosome, spliceosome, glycolysis /gluconeogenesis, RNA transport, oxidative phosphorylation, protein processing in the endoplasmic reticulum, MAPK signaling pathway - plant, methionine and cysteine metabolism, purine metabolism and RNA degradation. Unlike the previous study, this is for the first time, the present study identified these pathways may play key role in MYMIV resistance in *Vigna mungo.* Thus, information and transcriptomes data available in the present study make a significant contribution to understanding the genomic structure of *Vigna mungo*, enabling future analyses as well as downstream applications of gene expression, sequence evolution, and genome annotation.

## 1. Introduction

The *Vigna mungo* L (black gram) is one of the most important pulse crops of the Asiatic region (Kundu et al., 2019). As it is native to India, it has been widely cultivated since ancient times in South and South-East Asian countries (Patel et al. 2017). However, the fast industrialization and mechanization that are associated with shortening agricultural lands have severely affected crops developments via enhancing both abiotic and biotic stresses on crops plant (Gupta et al., 2018; Gouda et al., 2019; Donde et al., 2019; Gupta et al., 2019). Earlier studies have reported that pathogens, for instance, bacteria, viruses, fungi, osmolytes, and nematodes, severely attack host crops for sustaining their life cycle as well as growth (Chakraborty and Basak, 2018). However, in response to these pathogen, plants induce multiple signalling pathways at transcriptional as well as translational level, including activation of cytosolic Ca2+, mitogen-activated protein kinases (MAPKs), reactive oxygen species (ROS), fortification of plant cell wall, various transcription factors and initiation of several pathogenesis-related (PR) gene products that in turn either inhibit the growth or penetration of pathogen into the plant (Gupta et al., 2018; Gouda et al., 2019; Donde et al., 2019; Gupta et al., 2019).

In comparison to abiotic stress, in *Vigna mungo*, productivity is severely hampered through various insects, pests, and pathogens. Out of all factors, yellow mosaic disease (YMD) has the most damaging effect. In some cases, the loss may count up to 100%. Yellow Mosaic Disease of *Vigna mungo* is caused via the begomovirus Mungbean MYMIV (Mungbean Yellow Mosaic Indian Virus) (Patel et al. 2017). It belongs to geminiviridae family and is transferred via *Bemisia tabaci* (Ganguli et al., 2016). To date, several studies have been performed to understand the underlying biological processes associated with the response of *V. mungo* against MYMIV (Chakraborty and Basak, 2018; Ganguli et al., 2016). In 2017, Patel and the team reported that during MYMIV infection in *Vigna mungo*, mitogen-activated protein kinases (MAPK) gets activated, which in turn transduce signals of pathogen invasion to downstream molecules thereby causing diverse immune responses (Patel et al. 2017). Recently, Kundu and the team performed a study to understand the molecular basis of the MYMIV-stress response in *V. mungo* and identified transcriptomic differences amongst the susceptible (T9) and resistant (VM84) genotypes. Additionally, they also identified numerous plausible transcripts that may be associated with the *V. mungo* resistance against MYMIV. They also detected dissimilar gene expression in T9 and VM84 with 1679 and 2158 differentially expressed genes (DEGs), respectively. Genome-wide transcriptomic analysis in VM84 reflects a quick and strong immune reaction thereby suggesting efficient pathogen surveillance leading to activation of basal as well as induced immune responses. They also identified several functional pathways, mainly “plant hormone signal transduction” (ko04075), “starch and sucrose metabolism” (ko00500), “plant-pathogen interaction” (ko04626), “biosynthesis of secondary metabolites” (ko01110) and metabolic pathways” (ko01100) that gets upregulated during MYMIV-stress response in *V. mungo* (Kundu et al., 2019).

However, irrespective of all significant findings, Kundu and the team (Kundu et al., 2019) did not report about the functional annotation of the complete transcriptomes generated. Nevertheless, for identifying differential expressed transcripts, Kundu and team employed DESeq (Wang et al., 2010) packages of R (Team, 2014). Earlier several studies have reported that, in comparison to DESeq, recently developed edgeR package is more powerful as well as more sensitive to outliers (Ortutay and Ortutay, 2016). Though the procedure employed in both DESeq and edgeR are almost similar, these two packages compute library size normalization factors differently. While DESeq uses a relative log expression technique, edgeR employs trimmed means techniques (Ortutay and Ortutay, 2016). Nevertheless, the main differences lie in the approaches to estimate typical as well as gene-wide dispersion. While edgeR smooths gene-wide dispersion toward the common dispersion and moderates the differences, DESeq maximizes it if the variability of a gene is larger than the common value. Hence, in comparison to DESeq, the edgeR approach is more powerful as well as more sensitive to outliers (Ortutay and Ortutay, 2016). In 2014, Zhang and team analyzed results obtained from qRT-PCR as well as microarrays and reported that, in comparison to DESeq, edgeR performs better in terms of identifying true positives with the default FDR setting (FDR<0.05) (Zhang et al., 2014).

Considering the above information, in the present study, authors made an attempt to generate novel transcriptome of *Vigna mungo* by re-analyzing publicly available RNA seq data (Ganguli et al., 2016) via computational approaches. Subsequently, quality assessment followed by functional annotation of the complete transcriptome dataset was performed. Further, identification of differentially expressed transcripts was performed to identify more significant key pathways that play important role toward the resistance of MYMIV in *Vigna mungo.* The new transcriptomes generated and characterized in the present study make a significant contribution to understand genomic structure of *Vigna mungo*, aiding future analyses and downstream applications of sequence evolution, gene expression, and genome annotation.

## 2. Material and methods

### 2.1 Data collection

Raw Illumina Hiseq2000 RNAseq data of MYMIV resistant (control) (Accession no: SRR2058996) and MYMIV infected (Accession no: SRR2060093) *Vigna mungo* was downloaded from NCBI short read archive (SRA) archive (https://www.ncbi.nlm.nih.gov/sra). Later, these two SRA files were split into respective paired-end FASTQ files with the “fastq-dump” command of the SRA Toolkit (Staff, 2011).

### 2.2 Quality control

Quality of each paired-end FASTQ file was examined employing AfterQC separately (Chen et al., 2017). AfterQC automatically discards bad reads, identifies and removes bubble effects of the sequencer, trims read at both tail and front, recognizes sequencing errors, and corrects them partly. In the end, AfterQC generates a FASTQ file with excellent quality reads only. These files will be referred to as clean FASTQ files henceforth (Chen et al., 2017).

### 2.3 Initial *de novo* transcriptomes assembly

Consequently, all clean paired-end FASTQ files were subjected to *in silico* normalization followed by denovo transcriptomes assembling procedure via Trinity v.2.8.5 assembler with default parameters (Grabherr et al., 2011). Trinity reconstructs transcriptomes from RNA-seq data by combining three self-governing software modules, namely, Chrysalis, Inchworm, & Butterfly. Trinity divides sequence data into several distinct de Bruijn graphs, each denoting transcriptional complexity at a given locus/gene, and then processes each graph independently for extracting full-length splicing isoforms as well as for teasing apart transcripts generated from paralogous genes (Haas et al., 2013).

### 2.4 Quality assessment of the transcriptomes

Unlike Kundu and team (Kundu et al., 2019), before discarding redundant as well as poorly constructed transcripts, initial quality assessment of the transcriptomes generate was performed. We computed completeness via investigating the number of input RNA-seq reads that were characterized in our *de novo* assembled transcriptomes, as per the pipeline recommended in the Trinity package (https://github.com/trinityrnaseq/trinityrnaseq/wiki). Read representation was computed via mapping the cleaned reads back to their corresponding assemblies with the help of Bowtie2 v2.2.6 (“–local”, “–no-unal”) (Carruthers et al., 2018).

### 2.5 Functional annotation of the transcriptomes

As no functional annotation was performed earlier in the context of these transcriptomes data, for the first time in the present study, the author made an attempt to do the same by classifying them into various sequence classes, namely, lncRNA, SSR, and protein-coding sequences. Further, functional annotation of these protein-coding sequences was performed, followed by identifying key genes/transcripts that play an important role in resistance of MYMIV in *Vigna mungo*.

#### 2.5.1 Identification of long non-coding RNA

RNAplonc (https://github.com/TatianneNegri/RNAplonc) was employed for identifying long non-coding RNA from the assembled transcript dataset. RNAplonc utilizes a classifier approach for detecting lncRNAs in plants from mRNA-based data (Negri et al., 2019). RNAplonc was developed and trained with mRNA and lncRNA dataset from five plant species, namely, cucumber, thale cress, soybean, Asian rice, and western balsam-poplar (Negri et al., 2019).

#### 2.5.2 Identification of simple sequence repeats

Simple sequence repeats (SSRs) or microsatellites are short and tandemly repeated di-, tri-, tetra-or pentanucleotide motifs (Tautz and Renz, 1984). In comparison with other molecular markers, SSRs are dispersed and abundant in almost all genomes with a high level of polymorphism. Hence, SSRs analysis is a widely employed versatile tool in evolutionary studies and plant breeding programs. Frequency and relative abundance of complete SSRs were performed on complete transcriptome dataset using a Perl script named MIcroSAtellite (http://pgrc.ipk-gatersleben.de/misa/misa.html). The MISA was set to identify SSRs by unit size (x) and the least number of repeats (y): 2/6, 3/5, 4/4, 5/4, 6/4 (x/y) distance between two SSR sequences ≥100 bp (Han et al., 2018).

#### 2.5.3 Identification and removal of redundant protein-coding sequences

Subsequently, TransDecoder v3.0.0 (http://transdecoder.github.io/) was utilized for identifying and extracting all possible single best open reading frame (ORF) coding regions per transcript present in the assembled transcripts (Carruthers et al., 2018). Transcripts having ORFs < 200 bp length were further discarded via seqtk (https://github.com/lh3/seqtk). Later, redundant sequences were removed via clustering highly similar sequences with the help of CD-Hit v4.6.6 (https://github.com/weizhongli/cdhit/wiki) and keeping amino acid sequence identity as 1.00 (Carruthers et al., 2018). These protein sequences will be utilized further for scanning their homologs in three species of *Vigna*, i.e. *angularis, radiate* and *unguiculata.*

##### 2.5.3.1 Homology Search

For determining the degree of accuracy how assembled protein sequences were re-constructed to full- or near full-length, the non-redundant protein sequences were subjected to BLASTP searches (evalue=1e-3) against in-house protein database derived from protein sequence of genus *Vigna*. In-house database was developed with the option -makeblastdb present in the BLAST software. The protein sequence of *Vigna angularis* and *Vigna radiata* was obtained from the NCBI database (https://www.ncbi.nlm.nih.gov/protein) while that of *Vigna unguiculata* was obtained from the Phytozome database (https://phytozome.jgi.doe.gov/pz/portal.html).

##### 2.5.3.2 Functional annotation

Subsequently, functional annotation of each protein sequence was performed to detect gene ontologies (molecular function, biological pathway, and cellular component) and key pathways associated with them via various online tools, like BlastKOALA (https://www.kegg.jp/blastkoala/) and NetGO (http://issubmission.sjtu.edu.cn/netgo/). Further, in house program and manual inspection was performed to scan plant-specific gene ontologies and pathways. Signal peptide and transmembrane protein were detected through the Phobius (http://phobius.sbc.su.se/) tool. Protein domains present in each protein were detected via HMMER v.3.2,1 (https://github.com/EddyRivasLab/hmmer) against the Pfam database. Finally, the complete result was summarized together in excel sheet.

### 2.6 Transcript Abundance and Analysis of differentially expressed genes

Transcript abundance of each transcript present in the transcriptome file was performed through an alignment-based approach, namely Salmon (Patro et al., 2017), present in the Trinity package. Salmon quantify transcript abundance via mapping reads of each biological replicate against the respective assembled transcriptome. The alignment was performed via bowtie2. The mapping-based mode of Salmon works in two steps: (a) indexing and (b) quantification. The indexing phase works independent of the reads and is required to run only once on a specific reference transcripts set. The quantification step works the onset of RNA seq reads and is executed more frequently. After estimating transcript abundance for each condition, normalized expression values matrix combining both transcript quantification dataset was generated. Later this matrix was utilized to produce the expression level of each transcript via ExN50 analysis. Unlike Kundu and team, expression value file produced via Salmon was subjected to edgeR package of R to identify differential expressed transcripts in a disease condition. Transcripts having FDR□<□0.05, fold change >2 between resistant and infected samples were considered as key candidate genes.

### 2.7 Functional annotation and KEGG pathway enrichment analysis of differentially expressed genes

GO and pathway enrichment analysis of up-regulated and down-regulated DEGs was performed, separately, through an in-house program against the annotation file generated above.

## 3. Result And Discussion

### 3.1 Quality control

In the present study, we present novel and high-quality protein-coding transcriptomes for *Vigna mungo*. Initial inspection via AfterQC reveals that FASTQ of resistant and infected plants comprised 37.870 and 53.486 million paired-end reads respectively. Further, quality control analysis via AfterQC removed ∼0.544% and ∼0.538% bad reads from resistance and infected FASTQ file respectively (**Table 1**).

**Table 1:** Summary statics of FASTQ generated via AfterQC.

| **Parameter** | **RESISTANCE** | **INFECTED** |
| --- | --- | --- |
| Sequencing: | 2*101 pair end | 2*101 pair end |
| Total reads: | 37.870 M | 53.486 M |
| Filtered out reads: | 215980.000 (0.544%) | 301734.000 (0.538%) |
| Total bases: | 3.735 G | 5.275 G |
| Filtered out bases: | 382614442.000 (9.540%) | 537570743.000 (9.490%) |
| Estimated seq error: | 0.06% | 0.06% |
| Adapter trimmed reads: | 526.272 K | 515.589 K |
| Adapter trimmed bases: | 10.954 M | 10.241 M |
| Auto trimming | front:9, tail:0 | front:9, tail:0 |

### 3.2 Initial *de novo* transcriptomes assembly

The initial *de novo* transcriptomes assembly generated ∼183 MB transcriptomes file in FASTA format. Transcriptomes file is comprised of 146696 and 196647 number of genes and transcripts, respectively. Total GC% is 40.11. Total number of N50 for genes and transcripts are 1103 and 1566, respectively. N50 is the maximum length where at best 50% of the total assembled sequence lives in contigs of at least that length (Thunders et al., 2017). Median contig length of genes and transcripts is 397 and 499, respectively. Total assembled bases in genes and transcripts are estimated to be 103,122,525 and 178199140, respectively. Complete information about transcriptomic data is depicted in **Table 2**. The number of transcripts identified in the present study was ∼90000 higher than previous studies (Kundu et al., 2019; Singh et al., 2018). This might be due to more precise screening via AfterQC. However, no information about the number of genes was present in previous studies (Kundu et al., 2019; Singh et al., 2018).

**Table 2:** Summary of constituent data of Trinity-assembled *Vigna mungo* transcriptome.

|  |  |
| --- | --- |
| Total Trinity 'genes' | 146,696 |
| Total Trinity transcripts | 196,647 |
| Percent GC | 40.11 |
| Stats based on ALL transcript contigs |  |
| Contig N10 | 3844 |
| Contig N20 | 2922 |
| Contig N30 | 2345 |
| Contig N40 | 1932 |
| Contig N50 | 1566 |
| Median contig length | 499 |
| Average contig length | 906.19 |
| Total assembled bases | 178,199,140 |
| Stats based on ONLY LONGEST ISOFORM per 'GENE' |  |
| Contig N10 | 3356 |
| Contig N20 | 2445 |
| Contig N30 | 1895 |
| Contig N40 | 1467 |
| Contig N50 | 1103 |
| Median contig length | 397 |
| Average contig length | 702.97 |
| Total assembled bases | 103,122,525 |

**Table 3:**
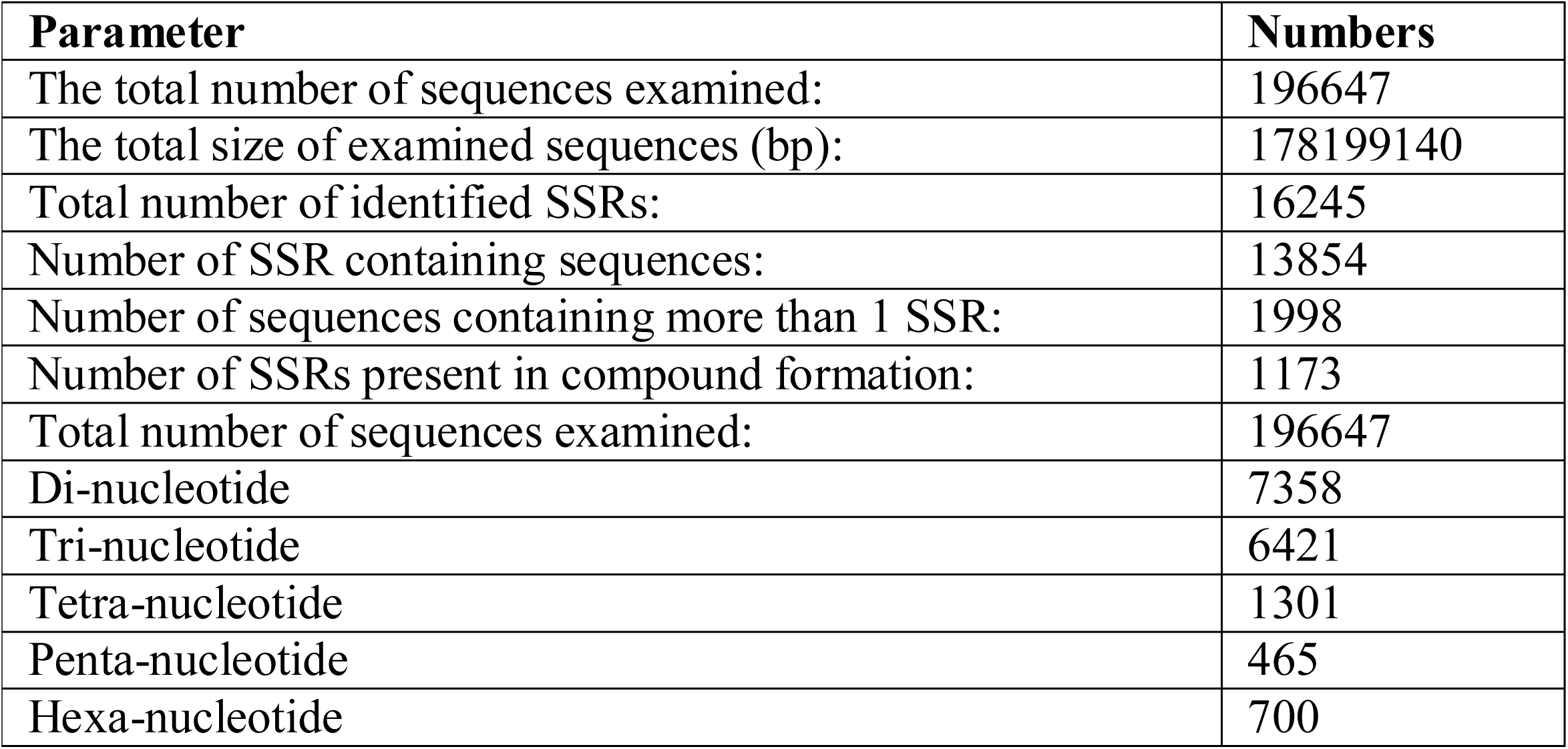
Summary of SSRs identified in the transcriptome of *Vigna mungo*.

| Parameter | Numbers |
| --- | --- |
| The total number of sequences examined: | 196647 |
| The total size of examined sequences (bp): | 178199140 |
| Total number of identified SSRs: | 16245 |
| Number of SSR containing sequences: | 13854 |
| Number of sequences containing more than 1 SSR: | 1998 |
| Number of SSRs present in compound formation: | 1173 |
| Total number of sequences examined: | 196647 |
| Di-nucleotide | 7358 |
| Tri-nucleotide | 6421 |
| Tetra-nucleotide | 1301 |
| Penta-nucleotide | 465 |
| Hexa-nucleotide | 700 |

### 3.3 Quality assessment of the transcriptomes

As an initial examination of the quality of the assembly, we mapped the generated transcriptome file back with the input reads files before filtering. Earlier studies have reported that more than 80% read mapping indicates good quality assembly (Haas et al., 2013). The obtained result reveals that 99.03% of the reads are successfully aligned. Thus, the generated transcriptome file was employed for downstream analysis.

### 3.4 Functional annotation of the transcriptomes

Identification and annotation of each sequence class performed separately.

#### 3.4.1 Identification of long non-coding RNA

In the present study, an analysis of the assembled transcript via RNAplonc identified 109126 lncRNAs in *Vigna mungo*. This accounts for 31% of the total transcripts (**Fig. 1A and Supplementary File 1**). Earlier, Singh and the team (Singh et al., 2018), employing the same raw FASTQ dataset, identified 2674 lncRNAs via the CPC Calculator tool (Kong et al., 2007). It is pertinent to note that 2256 lncRNAs from both studies share more than 80% of identity amongst themselves. In recent years, lncRNAs have got more attention because of their high involvement in gene expression regulation (Negri et al., 2019). lncRNAs are comprised of conserved domains that interact with various mRNAs, DNA, proteins, and miRNAs to carry out different biological functions. These lncRNAs along with several other molecules modulate numerous physiological processes, and disruption of these functions causes numerous diseases, including infectious diseases and cancer. These lncRNAs are responsible for modulating flux of genetic information (for instance transcription), chromosome structure regulation, the stability of messenger RNA (mRNA), splicing, and post-translational modifications (Fernandes et al., 2019).

**Fig. 1:**
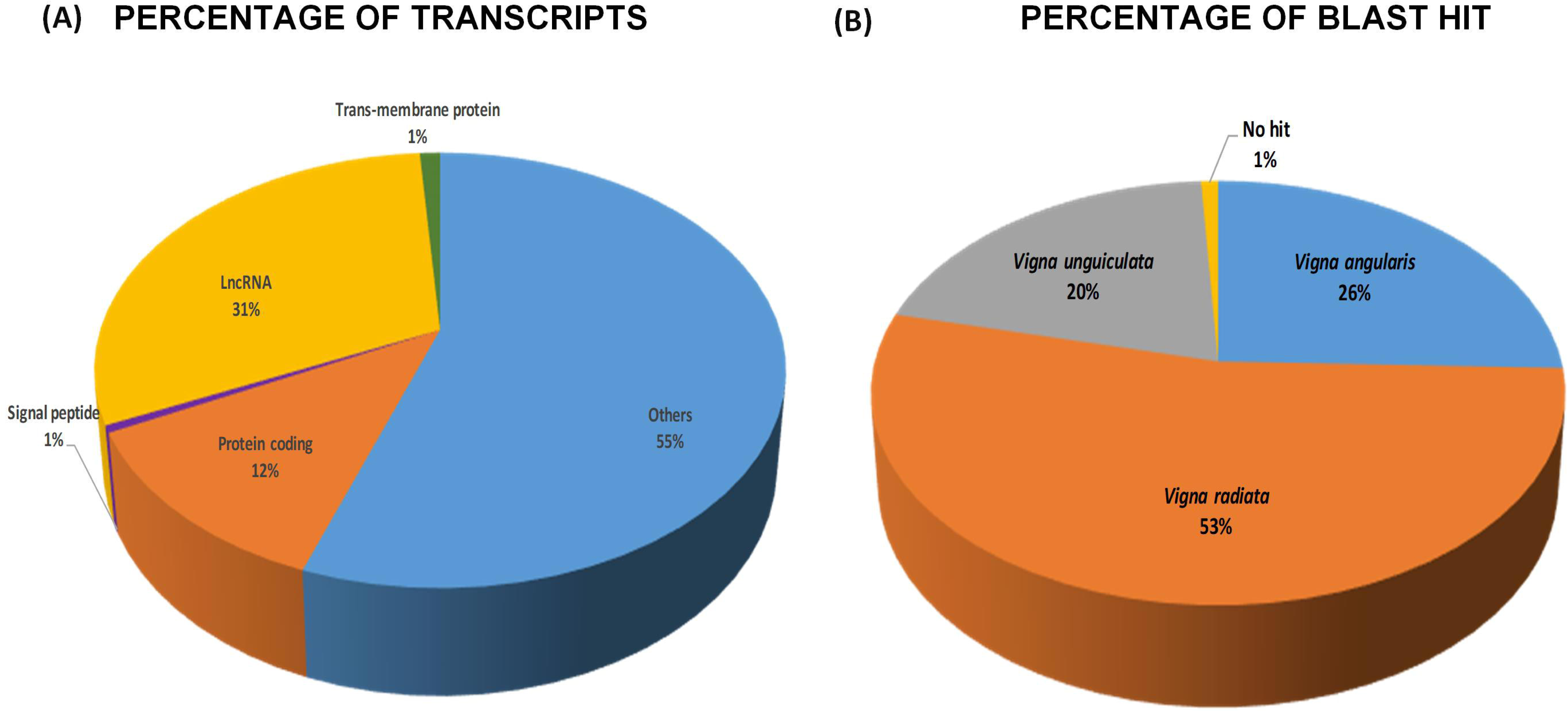
(A) Percentage of transcripts in each sequence class. (B) Percentage of BLAST hit for all unique 43194 protein-coding sequences against an in-house protein database.

#### 3.4.2 Identification of SSR molecular markers

As stated above, SSR is tandemly repeated short nucleotide units. As it successfully differentiates F1 and heterozygous populations, it is often referred to as a co-dominant marker (Brumlop et al. 2011). Till date, SSRs have been widely used in fingerprint accessions, variety identification, and diversity analysis, mapping of genes/QTL and marker-assisted selection (Sandhu 2013). SSRs markers are the first choice for crop improvement because of their locus specificity, reproducibility, and hypervariability in the genomes. The most important character is their codominant nature. However, the use of SSR molecular markers in molecular breeding has made it possible to assess the selection of diverse cultivars more efficiently for prospective utilization as parents (von Stackelberg et al., 2006). Till date, the complete analysis of SSR loci in the chloroplast or mitochondrial genomes has been done in rice (Rajendrakumar et al., 2007), bryophytes (Zhao et al., 2016), capsicum (Cheng et al., 2016), and soybean (Powell et al., 1995).

Earlier, SSR markers identified in closely related *Vigna* species were utilized for studying their transferability to *Vigna mungo*, and by employing those SSR markers, two linkage maps were also established in *Vigna mungo*. Nevertheless, the utility of transferable SSR markers in these linkage maps was restricted, and only 47 SSR loci were allotted to the 11 linkage groups. Hence, there is an urgent requirement for identifying and developing new polymorphic SSR markers in *Vigna mungo* (Souframanien and Reddy, 2015). In the present study, SSR analysis of all transcripts via MISA reported that out of 196647, 16245 SSR markers present in 13854 transcripts sequence. Top three repeat types were dinucleotide SSRs (7358), trinucleotide SSRs (6421), and tetra-nucleotide SSRs (1301), which altogether comprise ∼92% of the total SSRs. Amongst detected motif sequences, the top three most frequent SSRs were AG/CT (25.70%), AAG/CTT (11.94%), and AT/AT (11.65%) (**Table 2**). Earlier SSRs analysis of transcriptome in *Vigna mungo* reported that most frequent dinucleotide repeats are GA/TC (59.7%), GT/AC (19.5) and AT/TA (19.3%). Amongst the most frequent trinucleotides are GAA/CTT (25.6%) and GGT/ACC (14.7%) (Souframanien and Reddy, 2015).

#### 3.4.3 Identification and removal of redundant protein-coding sequences

Further identification of the protein-coding sequence present in the transcriptomes was performed through TransDecoder. At first, all possible 133613 ORFs were detected via “TransDecoder.LongOrfs” tool. Further, “TransDecoder.Predict” tool was employed to screen 85796 ‘single-best’ ORFs. Subsequent removal of ORFs having < 200 bp length via seqtk scanned 52286 ORF. Further, removal via CD-Hit v4.8.1 scanned 43194 ORFs. Thus, protein-coding sequences account for ∼14% of the total transcripts (**Fig. 1A**). As no functional annotation for these 43194 protein-coding sequences was performed earlier, for the first time in the present study, the author made an attempt to do the same. At first, sequence homology searches were perfumed against the in-house protein database of genus *Vigna*. Subsequently, functional annotation of each protein sequence was performed to detect gene ontologies and key pathways associated with them via various online tools, like BlastKOALA (https://www.kegg.jp/blastkoala/) and NetGO (http://issubmission.sjtu.edu.cn/netgo/).

##### 3.4.3.1 Homology Search

A BLASTP search of 43194 protein-coding sequences against in-house protein database revealed that, out of 43194 non–redundant transcripts, 53% matches with *Vigna radiate*, while 20% and 26% of the transcripts have a better match with *Vigna unguiculata* and *Vigna angularis*, respectively. This result is in accordance with earlier studies where authors have reported that phylogenetically *Vigna mungo* is more closely related to *Vigna radiate* than with *Vigna unguiculata* and *Vigna angularis* (Gupta et al., 2014; Vir et al., 2016; Asghar et al., 2018). However, 1% transcripts are unique to *Vigna mungo* and thus, show no hit with either of the three other species of *Vigna* (**Fig. 1B and Supplementary File 1**).

##### 3.4.3.2 Functional annotation of protein-coding sequences

Gene ontology and pathway enrichment analysis via NetGO and BlastKOALA, respectively reveals that these 43194 protein sequences are associated with 4682, 3871, 1147, and 191 unique biological processes, molecular function, cellular component, and pathways, respectively, in *Vigna mungo* (**Supplementary File 1)**. The top three important biological processes associated with these protein-coding sequences are cellular process and multicellular organismal process. Top three significant molecular functions associated with these protein-coding sequences are binding, heterocyclic compound binding, catalytic activity, protein binding, transferase activity, nucleic acid-protein binding, hydrolase activity, and catalytic activity. The top three important cellular components associated with these protein-coding sequences are organelle, intracellular part, and intracellular organelle part. The top three significant pathways associated with these protein-coding sequences are genetic information processing, carbohydrate metabolism, and environmental information processing (**Fig. 2**).

**Fig. 2.**
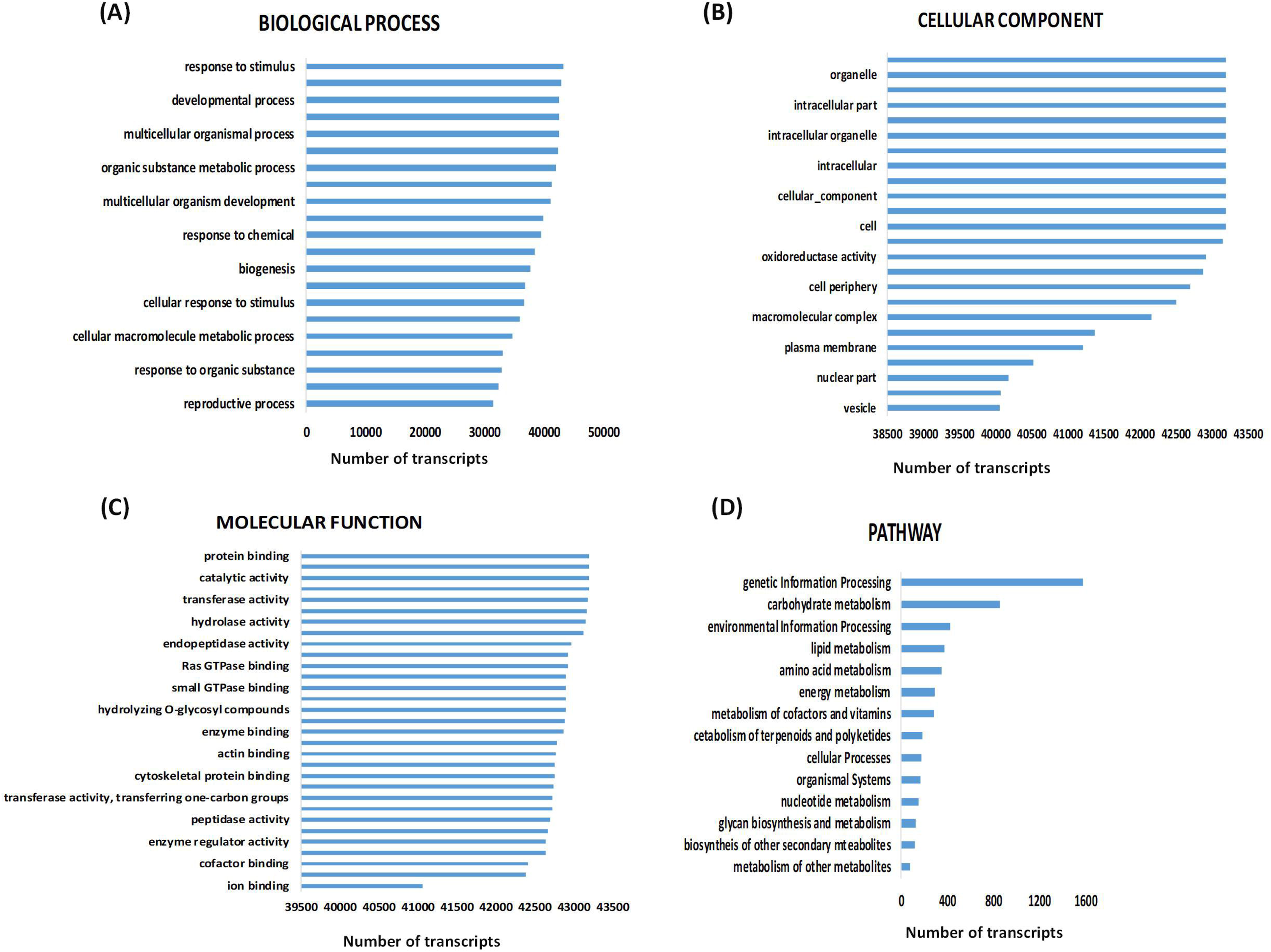
Most important gene ontology and pathways associated with protein coding sequences of *Vigna mungo*.

### 3.5 Transcript Abundance and analysis of differentially expressed genes

To estimate the level of transcript expression, at first, transcript quantification under both resistance and infection conditions was performed separately. Subsequently, normalized expression values matrix combining both transcript quantification dataset was generated. Later this matrix was utilized to produce the expression level of each transcript via ExN50 analysis. The obtained result revealed that, out of 196647, only 145715 transcripts are abundantly present in *Vigna mungo* under both resistant as well as infected conditions. Further, differential gene expression (DEGs) analysis between resistant and infection condition via edgeR package of R suggested that, out of 145715 DEGs, only 83344 have |logFc|>2 and FDR < 0.05. Out of 83344, 4536 and 78808 are significantly are down-regulated and up-regulated, respectively (**Supplementary File 2**). Thus, these 83344 transcripts play an important role in the resistance of MYMIV in *Vigna mungo*. It is pertinent to note that most of the transcripts get up-regulated in-response to MYMIV infection, which in turn helps *Vigna mungo* to sustain resistance against it (**Fig. 3)**. This is in accordance with original studies (Kundu et al., 2019), where authors have reported that 1242 and 437 are up-regulated and down-regulated during MYMIV virus infection in *Vigna mungo.* However, the number of DEGs identified in the present study is higher in comparison to the original studies, where Kundu and the team have identified only 2158 DEGs. This might be due to the more robust nature of the edgeR package of R (Ortutay and Ortutay, 2016).

**Fig. 3:**
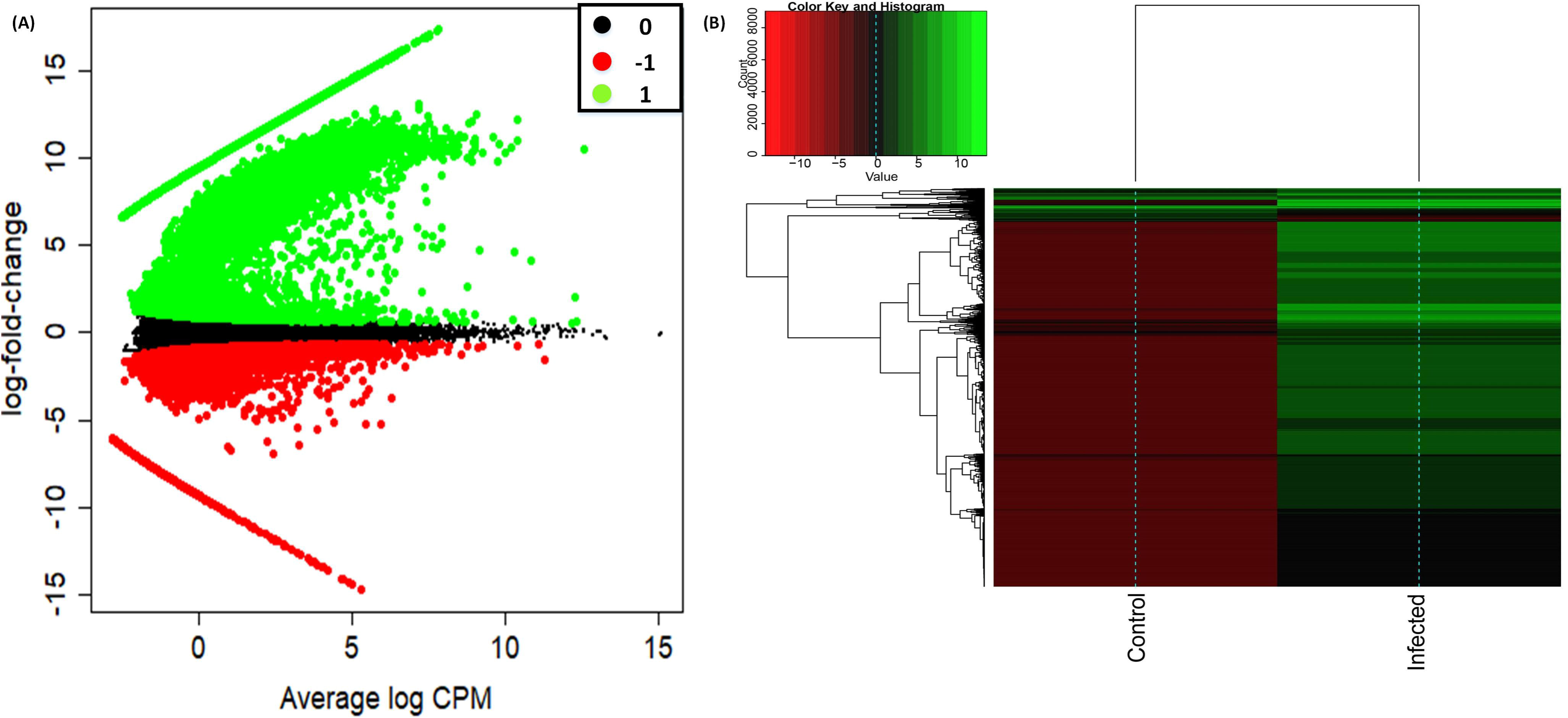
(A) A mean-difference plot plot representing the log-fold change and average abundance of each gene. Genes with fold-changes significantly greater than 2 are highlighted (Red=down-regulated, Green=up-regulated). (B) Heatmap diagram representing top 10000 differentials expressed genes (*p* < *0.01*) in transcriptome data of *Vigna mungo*. Heatmap clustering is based on transcripts abundance under both resistant and infected condition in *Vigna mungo*. Upregulated and downregulated genes are represented in green and red color respectively.

### 3.6 Functional annotation and KEGG pathway enrichment analysis of DEGs

Functional annotation and pathway enrichment analysis of up-regulated and down-regulated DEGs, separately, against annotation file, generate above, revealed that out 78808 up-regulated DEGs, 57551, 2100, 844, 1970 are lncRNA, protein, signal peptide, and transmembrane protein, respectively. Out of 4536 down-regulated, 4115, 88, 6, 5 are lncRNA, protein, signal peptide and transmembrane protein, respectively. This is for the first time we have reported that 61666 lncRNAs, 890 signal peptides, and 1975 transmembrane proteins that get differentially expressed during infection in *Vigna mungo*. Complete information about functional annotation and KEGG pathway enrichment analysis of DEGs is provided in **Supplementary File 2**. 2100 up-regulated protein modulates 120 unique pathways during infection in *Vigna mungo*. Out of 120, ten most important pathways are ribosome, spliceosome, glycolysis/gluconeogenesis, RNA transport, protein processing in the endoplasmic reticulum, oxidative phosphorylation, MAPK signaling pathway - plant, cysteine and methionine metabolism, purine metabolism and RNA degradation. Only eight unique pathways are modulated via down-regulated 88 down-regulated protein. Out of these 8, 5 are common to pathways of up-regulated DEGs. In original study Kundu and team have reported that 5 key pathways that play key role in MYMIV resistance in *Vigna mungo* are “plant hormone signal transduction” (ko04075), “starch and sucrose metabolism” (ko00500), “plant-pathogen interaction” (ko04626), “biosynthesis of secondary metabolites” (ko01110), metabolic pathways” (ko01100). This is for the first time, the present study identified spliceosome, glycolysis/gluconeogenesis, RNA transport, protein processing in the endoplasmic reticulum, methionine metabolism, and RNA degradation may also play key role in MYMIV resistance in *Vigna mungo.*

Earlier studies have reported that, in plant, defense against pathogens and insects are significantly modulated via Ribosome-Inactivating Proteins (RIPs). In 2018, Zhu and the team suggested that RIPs are toxic N-glycosidases that depurination eukaryotic as well as prokaryotic rRNAs, which in turn arrests protein synthesis during translation. RIPs are found in a wide range of plant species and have antibacterial, antifungal, insecticidal and antiviral, activities. However, how RIPs modulates these defenses against pathogens and insect pests is still a topic of debate (Zhu et al., 2018). Gluconeogenesis, a ubiquitous process, is associated with the production of glucose from a various non-carbohydrate carbon source, for instance, proteins and lipids. Glyceraldehyde-3-phosphate dehydrogenase (GAPDH), an important metabolic enzyme for energy production during glycolysis and gluconeogenesis, is also reported to play numerous cellular regulatory functions invertebrates as well as the plant (Henry et al., 2015). In 2015, Henry and the team reported that GAPDH knockouts *Arabidopsis* (KO lines) plant experiences increased disease resistance against *Pseudomonas syringae* pv. tomato. KO lines also exhibit increased programmed cell death, as well as enhanced electrolyte outflow in response to effector, triggered immunity (Henry et al., 2015).

Few studies have also reported about the relation between protein processing in the endoplasmic reticulum and plant resistance against the pathogen, the complete mechanism remains elusive to date (Farrokhi et al., 2008). Almost all proteins produced in the endoplasmic reticulum undergoes disulfide bond formation and protein folding during the oxidative folding process. Oxidative folding is modulated via numerous enzymes, including the family of protein disulfide isomerases (PDIs), as well as other proteins that supply oxidizing equivalents to PDI family proteins, like ER oxidoreductin 1 (Ero1) (Urade 2019). The MAPK signaling pathways is another most important pathway that plays key role during plant resistance against various biotic and abiotic stresses, as well as in response to plant hormones such as auxin, ethylene and abscisic acid. In 2019, Wan and the team reported that innate immunity in both animals and plants is activated via pattern recognition receptors (PRRs) in response to microbe-associated molecular patterns (MAMPs) for providing the first line of inducible defense (Wan et al., 2019). Plant receptor protein kinases (RPKs) represent the main plasma membrane PRRs perceiving diverse MAMPs. Additionally, RPKs also detect secondary danger-inducible plant peptides as well as cell-wall signals. Both forms of RPKs initiates fast as well as convergent downstream signaling networks modulated via mitogen-activated PK (MAPK) and calcium-activated PKs cascades. They also observed the integration of both long-term as well as late responses associated with MAPK and calcium-activated PKs cascades into the networks in innate immunity. In 2017, Patel and the team identified MAPKs homolog in the defense signaling pathway of MYMIV infected *Vigna mungo* (Patel et al., 2017). They also characterized a MAP kinase (VmMAPK1), which was induced upon MYMIV-inoculation in resistant *V. mungo*. VmMAPK1 protein contained in the nucleus as well as cytoplasm and possessed phosphorylation activity in *in vitro*. Phylogenetic analysis revealed that VmMAPK1 is closely related to other plants-stress-responsive MAPKs. The ability of VmMAPK1 to restrict MYMIV multiplication was validated by its ectopic expression in transgenic tobacco. This study suggested that overexpression of VmMAPK1 resulted in the considerable upregulation of defense-responsive marker PR genes. Both mRNA and protein of VmMAPK1 were accumulated upon MYMIV infection (Patel et al., 2017).

The cysteine and Methionine are also very essential in some plants because it improves the protein quality of some important legume crops (Joshi et al., 2019). Romero and the team reported that cysteine occupies central metabolic pathways in plants because it is a reduced sulfur donor molecule and is associated with the synthesis of defense compounds and essential biomolecules. Furthermore, cysteine plays important role in the redox signaling processes of various cellular compartments and is synthesized in sulphate assimilation pathway via the incorporation of sulphide to O-acetylserine, catalysed by O-acetylserine (thiol) lyase (OASTL). OASTLs are mainly situated in the cytosol, mitochondria, and chloroplasts, and is involved with complex array of isoforms and subcellular cysteine pools (Romero et al., 2014). Recently, Joshi and team reported that γ-Glutamyl-S-methylcysteine is present in Phaseolus and numerous Vigna species. γ-Glutamyl-S-ethenylcysteine, an antinutritional compound, is present in *Vicia narbonensis* (Joshi et al., 2019). Farrokhi et al. (2008) reported antimicrobial peptides are cysteine-rich basic peptides with distinct sequences (Farrokhi et al., 2008).

Nucleotide metabolism functions in all living organisms and represents an evolutionarily ancient and indispensable complex of metabolism pathways that are of supreme importance for plant metabolism and development (Zrenner et al., 2006). In plants, nucleotides can be synthesized *de novo* from 5-phosphoribosyl-1-pyrophosphate and simple molecules (e.g., CO2, amino acids, and tetrahydrofolate). In 2006, Zrenner and Team reported that nucleotides are degraded to simple metabolites molecules such as phosphate, nitrogen, and carbon into the central metabolic process (Zrenner et al., 2006). Despite extensive biochemical knowledge about purine and pyrimidine metabolism, comprehensive studies of the regulation of this metabolism in plants are only starting to emerge. Extracellular purine nucleotides capable of regulating plant development, defense and stress responses by acting in part as agonists of plasma membrane calcium channels. The Purine metabolism is stimulating by ATP release include wounding, osmotic stress and elicitors (Dark et al., 2011).

RNA degradation into the cell is one of the major metabolic processes that modulate gene downregulation. RNA degradation takes place in cells by several metabolic processes like RNAi, RISC CRIPER cas9 protein and RNA-DNA methylation of foreign molecules (Filippova et al. 2019). Recently, Muhammad and team suggested that RNA interference (RNAi) suppresses gene expression both transcriptionally and post-transcriptionally for resisting the virulence of pathogens by employing three distinct groups of proteins. For the first time, RNAis was discovered in early 1990 and proposed that introduced genes plausibly co-suppressed the endogenous genes. However, later it was observed that homologous RNA sequences were the main reason for suppressing internal genes (Muhammad et al., 2019). Initially, RNA silencing was employed in that study of animals. However, because of the presence of an extensive amount of pathogens and their threats to plants, later RNA silencing was also employed in crop research (Muhammad et al., 2019). RNA silencing mechanism is reported to initiated with the generation of ∼20 nucleotides small RNA (sRNAs), which in turn lead to the production of several other key products, namely, RNA-dependent RNA polymerase (RDRs), Dicer-like protein (DCL) and Argonaute (AGO) protein. The DCL proteins produces small RNAs (sRNAs) from a double stranded RNAs (dsRNAs) precursor and subsequently integrates it into RNA-induced silencing complexes (RISCs). Based on their origin as well as formation, these sRNAs are further classified as siRNAs or microRNAs (miRNAs). AGO proteins carry out most of the RISCs function, bind the sRNAs and interact with homologous RNAs, which in turn influences endonuclease activity, DNA methylation or translational repression of mRNAs. RDR enzymes is associated with dsRNAs synthesis employing single-stranded RNAs (ssRNAs) as the templates, which are subsequently processed further via Dicer-like (DCLs) proteins and start a new round of RNA silencing (Muhammad et al., 2019). Recently, Wang and the team reported that microRNAs that are associated with the RISCs play a significant role in plant development, either via obstructing translation or via targeting mRNA for cleavage. For few years, the list of numerous miRNAs, their confirmed targets as well as developmental effects has expanded, as has the realization that they are highly conserved throughout evolution and that small RNAs can play a direct role in cell-cell signaling (Wang et al., 2019). However, to the best of our knowledge, this is for the first time we report about the involvement of oxidative phosphorylation and spliceosome in the plant resistance against biotic or abiotic stress. Thus, information obtained from the present study will be highly useful for understanding mechanisms associated with MYMIV-resistance in *Vigna mungo*, which in turn will be highly helpful in controlling MYMIV-infection, thereby increasing yield in *Vigna mungo*.

## 4. Conclusion

Owing to the lack of reference genome, authors made an attempt to release new transcriptome, lncRNA, and protein-coding sequences of *Vigna mungo*, including SSR markers. As quality assessment of the generated transcriptomes suggested good quality, transcriptomes generated in the present study provide more robust resources for future genomic investigation in *Vigna mungo*. Additionally, this also provides vital tools, which, in near future, can be employed for performing comparative studies in *Vigna* species of evolutionary, ecological and economic interest. Thus, the new transcriptomes generated and characterized in the present study make a significant contribution to understand the genomic structure of *Vigna mungo*, facilitating future analyses and downstream applications of sequence evolution, genome annotation, and gene expression.

## Acknowledgement

Authors would like to thank Mr. Jan Benzenberg, Incident Manager, Carrier Radio Access Networks (Germany), for providing the computational facility to carry out a functional annotation.

## Data Availability

Transcriptome, lncRNA, protein sequence in FASTA format and gene ontology of complete protein-coding sequences in .xlsx format is available at Github (https://github.com/Manojbioinfo/Vigna).

